# A highly stringent high-throughput screening assay for identifying transcription-modulating agents of HIV-1 latency

**DOI:** 10.64898/2026.01.07.698296

**Authors:** Chhavi Saini, Agnita Roychowdhury, Shruti Gaikwad, Yuvrajsinh Gohil, Arun Panchapakesan, Payel Das, Saravanan Thiyagarajan, Murali Ramachandra, Susanta Samajdar, Ravi Manjithaya, Udaykumar Ranga

**Affiliations:** HIV-AIDS Laboratory, Molecular Biology and Genetics Unit, Jawaharlal Nehru Centre for Advanced Scientific Research, Jakkur, Bengaluru, Karnataka, India; Molecular Biology Laboratory, Y. R. Gaitonde Centre for AIDS Research and Education (YRG CARE), Chennai, Tamil Nadu, India; Aurigene Oncology Ltd, Electronic City Phase I, Bengaluru, Karnataka, India; Autophagy Laboratory, Molecular Biology and Genetics Unit, Jawaharlal Nehru Centre for Advanced Scientific Research, Jakkur, Bengaluru, Karnataka, India; Institute of Laboratory Medicine, Clinical Chemistry and Molecular Diagnostic, University Hospital, Leipzig, Germany

**Keywords:** HIV-1, high-throughput screening, LTR variants, latency-modulating agents, dual-reporter cell line

## Abstract

HIV-1 latency remains a central obstacle to curing infection, and current latency-modulating agents (LMAs) suffer from poor specificity and inconsistent efficacy. To enable discovery of small molecules (SMs) that directly target the viral master transcriptional regulatory circuit (MTRC), we developed a highly stringent, dual-reporter high-throughput screening (HTS) assay based on a natural HIV-1 subtype C long terminal repeat (LTR) variant, LRhR-HC, which exhibits markedly reduced transcriptional noise and a high activation threshold. We engineered Jurkat cells to stably harbour two independent reporter cassettes driven by the tough-to-activate LRhR-HC-LTR and the canonical LR-HHC-LTR, enabling simultaneous detection of latency-promoting and latency-reversing activities through both fluorescent and secreted enzymatic reporters. This dual-reporter line responded robustly and predictably to conventional latency-reversal agents (LRAs) and latency-promoting agents (LPAs), validating assay responsiveness. Z′-factor measurements demonstrated excellent assay performance, with values ranging from approximately 0.7 across different formats, confirming a strong dynamic range and reproducibility. Screening of the 1,520-compound Prestwick chemical library (PCL), which includes FDA-approved drugs, identified 27 candidate LPAs, including known agents such as Spironolactone and Aminacrine, thereby validating the assay specificity while revealing several novel inhibitory molecules. A mechanistically focused panel of 10 additional compounds yielded two putative LPAs and one LRA, with secondary analyses confirming their latency-modulating activities and cytotoxicity profiles. Collectively, this HIV-1C-derived dual-reporter platform provides a stringent and flexible HTS system for identifying LMAs with potential applications in both ‘block-and-lock’ and ‘shock-and-kill’ cure strategies.

## I. Introduction

Since the identification of Human Immunodeficiency Virus type 1 (HIV-1) in the 1980s, substantial effort has gone into developing effective therapies (1, 2). Antiretroviral Therapy (ART) suppresses viral replication to undetectable levels in people living with HIV (PLWH) (3, 4), but it does not eliminate latent proviruses (5, 6). These persist in resting CD4^+^ T cells and other lineages (7) and cause rapid viral rebound if ART is stopped, necessitating lifelong treatment. Within the memory CD4^+^ T cells, central memory cells (T_CM_) form a primary latent reservoir due to their long lifespan and low turnover. In contrast, in PLWH with low CD4^+^ T cell counts, transitional memory cells (T_TM_) become the dominant reservoir and persist through homeostatic proliferation (8, 9). Most latent proviruses are defective, with only a minority being intact and inducible. Inducibility varies by integration site (10–13), and the intact inducible reservoir shows minimal decay even during long-term ART (7).

ART is associated with several adverse effects, although recent improvements have made long-term regimens far more tolerable, including reductions in mitochondrial toxicity (14). Nonetheless, patients may still experience gastrointestinal symptoms, metabolic disturbances, weight gain, bone mineral density loss, renal or hepatic abnormalities, and other drug-specific toxicities (15, 16). Another limitation of ART is the emergence of drug-resistant viral variants, which are enriched under the selective pressure of the therapy (17). These challenges highlight the need for therapeutic strategies that directly target the latent HIV-1 reservoir.

Two main therapeutic strategies using latency-modulating agents (LMAs) are under investigation to address the latent HIV reservoir. The first, ‘shock-and-kill’, aims to reactivate latent proviruses with latency-reversal agents (LRAs). Several classes of LRAs have been identified, including PKC agonists, epigenetic modifiers, Akt pathway activators, SMAC mimetics, and MAPK agonists (18–20). The alternative ‘block-and-lock’ strategy seeks to drive the virus into a deeper, irreversible latent state using latency-promoting agents (LPAs). Only a few LPAs are known, with didehydro-cortistatin A (dCA) being the most potent, which demonstrated reduced viral rebound after ART interruption, including in a macaque model (21, 22).

To address the need for new LMAs, we developed a high-throughput screening (HTS) assay using a natural HIV-1 subtype C (HIV-1C) LTR variant, LRhR-HC. The canonical HIV-1C LTR (referred to here as LR-HHC) contains one LEF-1α/TCFα (L) and one RBEIII (R) binding site in the viral modulator region, and three NF-κB motifs (two variants, H and C) in the enhancer region (23). The LRhR-HC variant differs from LR-HHC in two key ways. First, its modulator region contains one L site but two R sites, separated by an additional NF-κB variant motif (h) (**Figure 1A**). Second, its enhancer contains only two NF-κB motifs (H and C) (23). Thus, compared with LR-HHC, the variant LTR gains an additional RBEIII site but possibly loses one functional NF-κB element. These compositional differences make the LRhR-HC promoter substantially more resistant to reactivation, requiring a markedly higher activation threshold for latency reversal than the canonical LR-HHC promoter (24, 25). Accordingly, we designate the conventional LR-HHC as a ‘simple-to-activate’ LTR and the variant LRhR-HC as a ‘tough-to-activate’ LTR.

**Figure 1.**
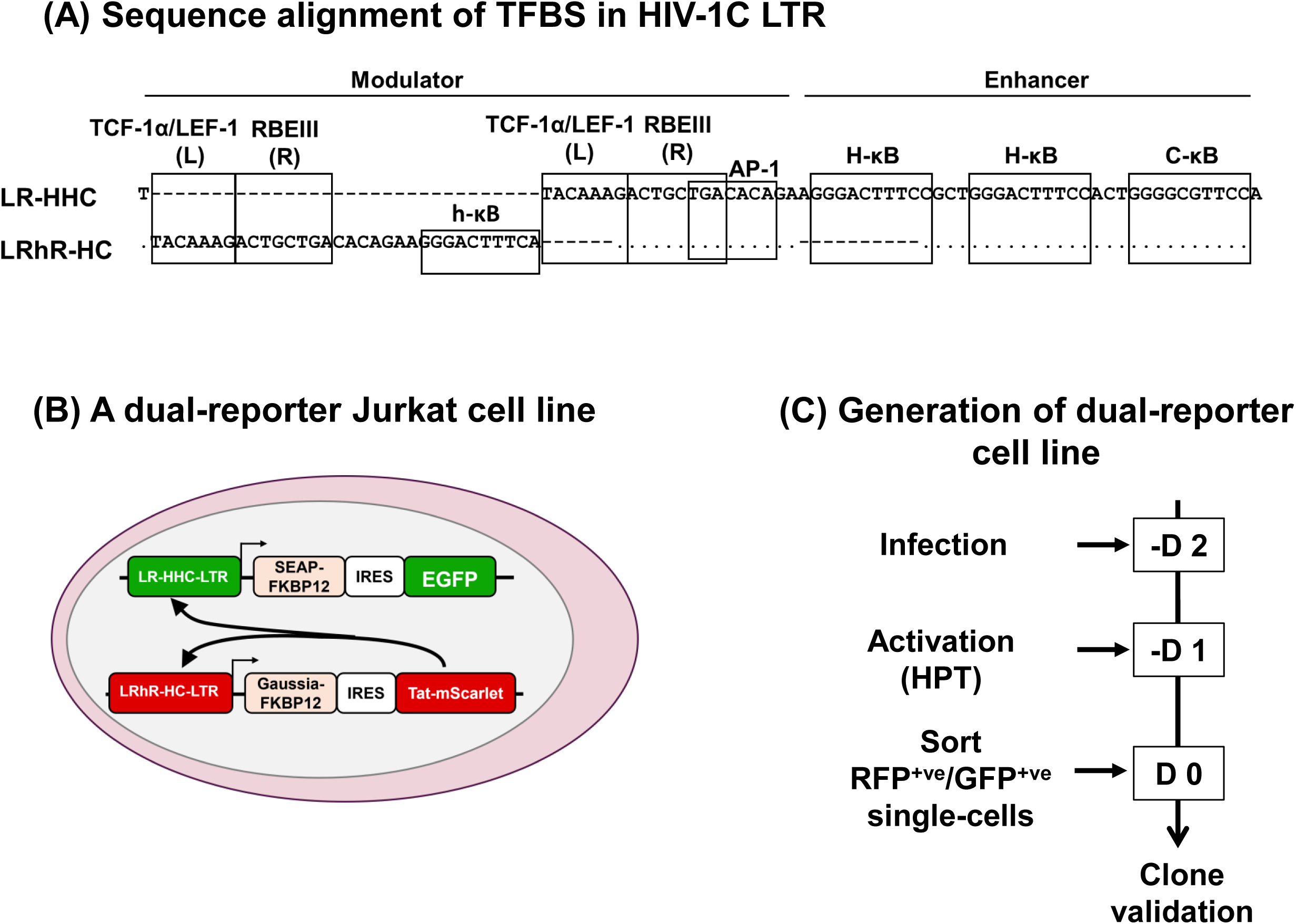
Generation of HTS dual-reporter Jurkat cell line. (**A**) HIV-1C LTR sequence alignment depicting the transcription factor binding sites (TFBS) variations in LR-HHC vs. LRhR-HC LTRs. Dots and dashes represent sequence homology or deletion, respectively. The modulator and enhancer regions of LTR are compared, showing TCF-1α/LEF binding sites (L), RBEIII sites (R), NF-κB sites (H/C-κB), κB-like site (h-κB), and AP-1 site, each separated by a box. (**B**) The dual-reporter Jurkat cell line harbours two different sub-genomic and reporter viral strains, LR-HHC and LRhR-HC, each characterised with differential activation threshold requirements. The simple-to-activate LR-HHC-LTR co-expresses a secretory alkaline phosphatase (SEAP) and an enhanced green fluorescent protein (EGFP) under the control of an internal ribosome entry site (IRES) element. The tough-to-activate LRhR-HC-LTR co-expression of the Gaussia luciferase and a Tat-mScarlet fusion protein under the IRES element. Secretory enzymes are tagged with an FKBP12 (FK506- and rapamycin-binding protein) destabilisation domain. Note that the expression of Tat is controlled by the more stringent LRhR-HC-LTR. (**C**) An experimental schematic representing a timeline in days (D) for generating the dual-reporter cell line for the HTS assay.

Conventional LMA-screening assays typically use the canonical HIV-1 LTR to drive a reporter gene expression (26–29). However, several intrinsic features of the HIV-1 LTR make it poorly suited for high-throughput identification of small-molecule inhibitors (SMIs) to target the viral master transcriptional regulatory circuit (MTRC) (30). First, Tat-independent transcription relies heavily on NF-κB-mediated signaling (31). Second, the Tat-dependent and Tat-independent transcriptional activities exclusively use the host transcription machinery; thus, except for Tat, all the other factors regulating viral transcription are derived from the host (32, 33). Consequently, many hits generated using the canonical LTR lack specificity because they target general host transcription pathways rather than the viral promoter itself. A third major limitation is the unusually high transcriptional noise produced by the HIV-1 LTR relative to cellular promoters (34, 35), a property thought to facilitate switching between active and latent states (36). This elevated background greatly increases false-positive rates in HTS assays based on the canonical LTR. In contrast, the LRhR-HC-LTR displays significantly low-level background transcriptional noise (37). Therefore, HTS assays built on this variant LTR are likely to reduce the background noise of the screening.

Here, we present a Jurkat cell-based dual-reporter HTS assay that leverages the low background-noise characteristics of the LRhR-HC-LTR. Using this platform, we screened the FDA-approved Prestwick chemical library (PCL) and identified several promising hits.

## II. Results

### Generation and selection of a dual-reporter Jurkat cell clone for high-throughput screening

Jurkat reporter cells harbour two different sub-genomic, reporter viral strains featuring LTRs of contrasting activation threshold requirements. Each LTR co-expresses a secretory protein and a fluorescent protein, allowing for independent, differential, and specific monitoring of the transcriptional activity of each promoter (**Figure 1B**). First, Jurkat cells were infected with a viral strain containing the tough-to-activate HIV-1C LRhR-HC-LTR, which co-expressed Gaussia luciferase (GLuc) and Tat-mScarlet (RFP) fusion protein under the control of an IRES element. Following infection, RFP^+ve^ cells were subjected to single-cell sorting, and the resulting cell clones were characterized for reporter gene expression. Subsequently, the intermediate cell clone, LRhR-HC-GLuc-IRES-Tat-mScarlet, was super-infected with a viral strain that co-expressed Secretory Alkaline Phosphatase (SEAP), followed by EGFP controlled by the IRES element under the control of the simple-to-activate LR-HHC-LTR (**Figure 1C**). The super-infected cells were activated with the HPT activator cocktail (5 mM HMBA, 5 ng/mL PMA, and 10 ng/mL TNF-α) for 24 hours, and EGFP/RFP dual-positive cells were single-cell sorted. Of the approximately forty dual-positive single-cell clones, four clones (9, 14, 19, and 39) that demonstrated superior proliferation were selected for further characterization.

Next, the four selected subclones (9, 14, 19, and 39) were activated using optimal or sub-optimal concentrations of the HPT activator cocktail, and reporter gene expression was quantified at 24 hours using either flow cytometry or enzymatic luminometry. Under the conditions of optimal activation, the expression levels of the two fluorescent proteins, EGFP (LR-HHC) and mScarlet (LRhR-HC), were comparable, regardless of the activation threshold requirements of the LTRs (**Figure 2A**, top panel). At a lower level of activation, the fluorescent protein expression was proportionately low; however, as expected, the tough-to-activate LRhR-HC-LTR showed a significantly lower level of gene expression than the canonical LR-HHC-LTR. Thus, the overall gene expression from the reporter cells demonstrated a dose-dependent response profile. Under all experimental conditions, cell viability remained high (**Figure 2A**, bottom panel).

**Figure 2.**
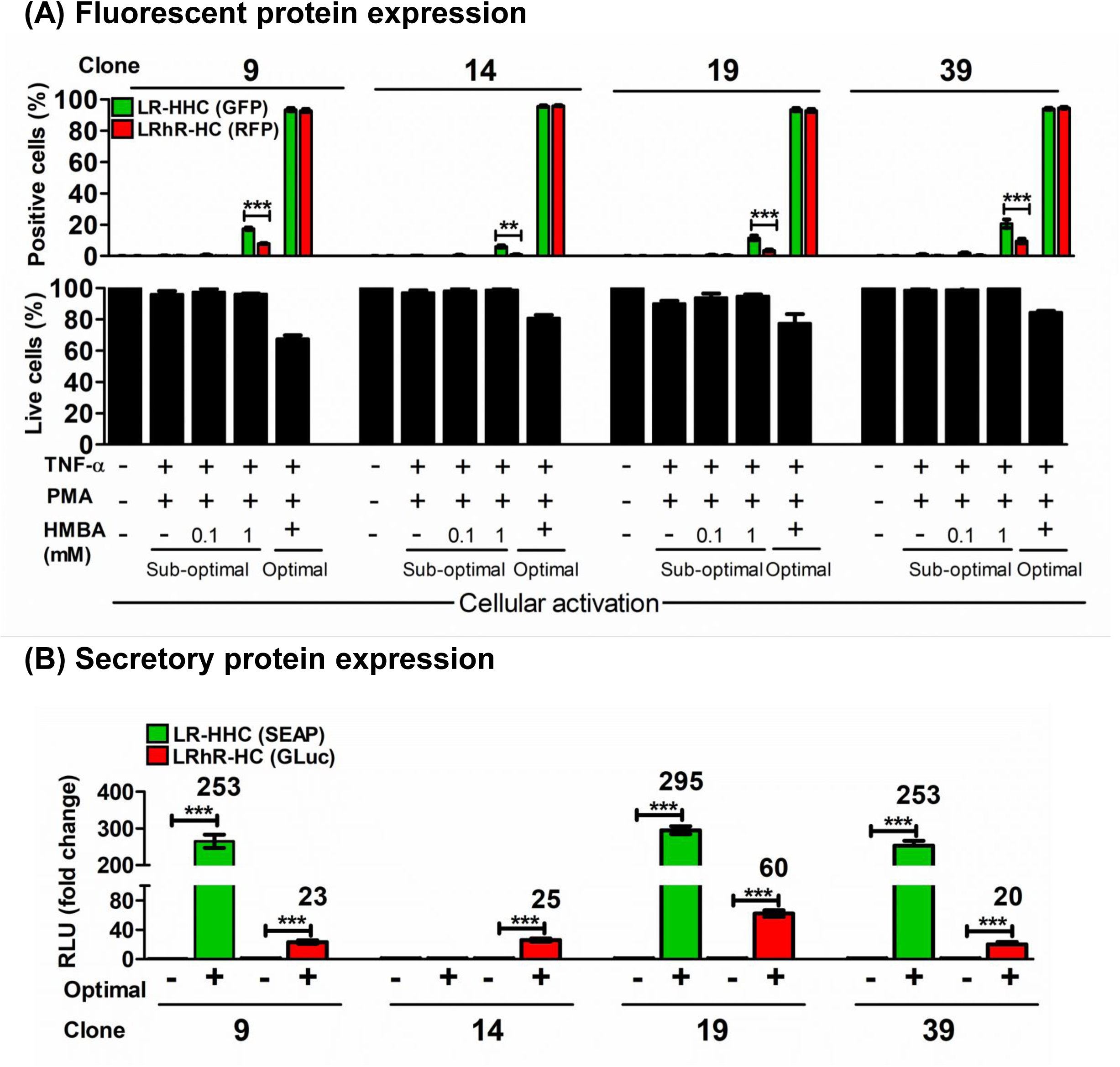
Clonal selection of the dual-reporter HTS Jurkat cell line. Following single-cell sorting, approximately 40 dual-positive Jurkat clones were obtained. Four clones (No. 9, 14, 19, and 39) were selected for further validation based on the activation profiles of both the LTRs (data not shown). (**A**) GFP or RFP co-expression following cellular activation. Cells were subjected to the HPT cocktail, a mix of three cellular activators, at an optimal (5 mM HMBA, 5 ng/ml PMA and 10 ng/ml TNF-α) or sub-optimal (0.1 or 1 mM HMBA, 1.5 ng/ml PMA and 0.625 ng/ml TNF-α) activation conditions for 24 hr. Cell viability was also monitored under all conditions (lower panel). (**B**) Co-expression of secretory enzymes following HPT activation (+), or no activation (-). Fold change in the reporter gene expression is depicted above each plot. Data plotted in triplicates, mean ± SD, statistical test – Student’s t-test, **p < 0.01, ***p < 0.001. Clone 19 was selected for further experimentation based on the reporter gene expression profile.

The expression levels of the secretory proteins – SEAP (LR-HHC), and GLuc (LRhR-HC) - were also assessed under conditions of optimal HPT activation. Three of the four clones (9, 19, and 39) expressed both enzymes at significantly higher levels following activation, although clone 14 failed to express SEAP (**Figure 2B**). For example, clone 9 expressed SEAP and GLuc at 253- and 23-fold higher levels, respectively. Following this analysis, we selected clone 19, hereafter referred to as the reporter cell line, due to its higher fold activation of GLuc. Thus, we successfully engineered a dual-reporter cell line that co-expresses two different fluorescent proteins and two secretory enzymes in response to cell activation. Importantly, the expression levels of these reporter proteins are differentially regulated by the activation threshold differences of two natural HIV-1 LTRs. Notably, the reporter expression under the tough-to-activate LRhR-HC-LTR provides a highly stringent screening assay, which is expected to be refractory to the cellular background transcription noise. Furthermore, while the expression of the fluorescent proteins can enable periodic clonal selection via sorting, that of the secretory enzymes can serve as a primary readout in the HTS assay. To maintain the additional stringency during the HTS assay, both secretory proteins – SEAP and GLuc are tagged with a degradation domain – FKBP12 degradation domain (FK506- and rapamycin-binding protein), which reduces their half-life to 2 hours (38). Of note, in several other experimental formats (**Figure 3-6**), we employed a Flash Gaussia luciferase substrate, which yields significantly higher luciferase readings.

**Figure 3.**
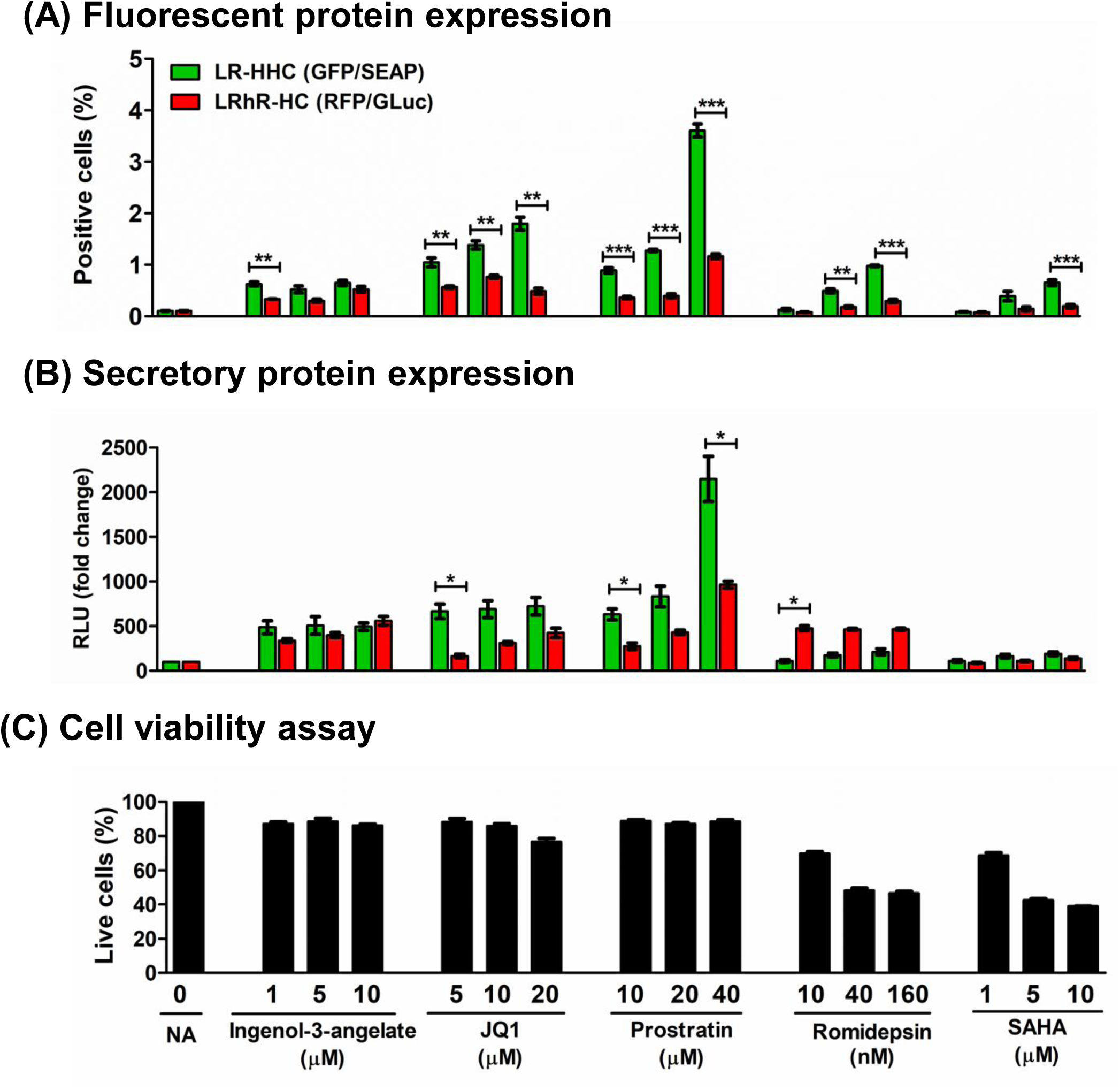
Latency-reversing agents can activate reporter gene expression from both LTRs. A few conventional LRAs were evaluated at varying concentrations for the activation of both LTRs for 24 hr. The expression profiles of the respective (**A**) fluorescent proteins and (**B**) secretory enzymes were monitored. (**C**) Cell viability was monitored in all conditions. Data plotted in triplicate, mean ± SD, statistical test – Student’s t-test, *p < 0.05, **p < 0.01, ***p < 0.001.

### The dual reporter cells are responsive to conventional LMAs

We validated the suitability of the engineered reporter clone against known LRAs and LPAs. The cells were exposed to varying concentrations of five conventional LRAs - Ingenol-3-angelate (39), JQ1 (30), Prostratin (40), Romidepsin (41), and SAHA (42), followed by quantification of secretory and fluorescent proteins. The LRAs successfully reactivated the two latent proviruses in the reporter cells; however, to varying levels (**Figure 3A**). Prostratin demonstrated the highest levels of viral reactivation among all LRAs. For instance, at a 40 µM concentration, Prostratin reactivated 3.6 ± 0.2% and 1.2 ± 0.1% of EGFP^+ve^ and RFP^+ve^ cells, respectively, as determined by flow cytometry. A similar trend was observed for the secretory reporter proteins, with 2,048.3-fold and 880.1-fold increases in SEAP (LR-HHC) and GLuc (LRhR-HC) expression, respectively, relative to the no activation control (**Figure 3B**). Cell viability was monitored under all conditions where maximal cell death was caused by Romidepsin and SAHA, with minimal LTR activation (**Figure 3C)**.

Next, we evaluated the effects of three known LPAs – Aminacrine (43), Apigenin (44), and Spironolactone (45). Cells were exposed to a sub-optimal HPT activator cocktail and three increasing concentrations of each LPA. All the LPAs exhibited a dose-dependent suppressive effect on both the LTRs (**Figure 4**). For example, treatment with Spironolactone at 20 µM reduced the activation of EGFP^+ve^ cells from 14.1 ± 1.4% to 0.16 ± 0.07%, and similarly, RFP^+ve^ cells decreased from 8.3 ± 0.9% to 0.12 ± 0.07% (**Figure 4A**). A similar trend was observed for secretory proteins, with Spironolactone reducing SEAP expression by 10.1-fold and for GLuc by 28.4-fold compared to the sub-optimal activation control (**Figure 4B**). Cell viability was also monitored under all conditions, and only minimal cytotoxicity was observed (**Figure 4C**). In summary, the validation experiments demonstrated that the dual-reporter cell line is responsive to conventional LRAs and LPAs in a manner consistent with expectations.

**Figure 4.**
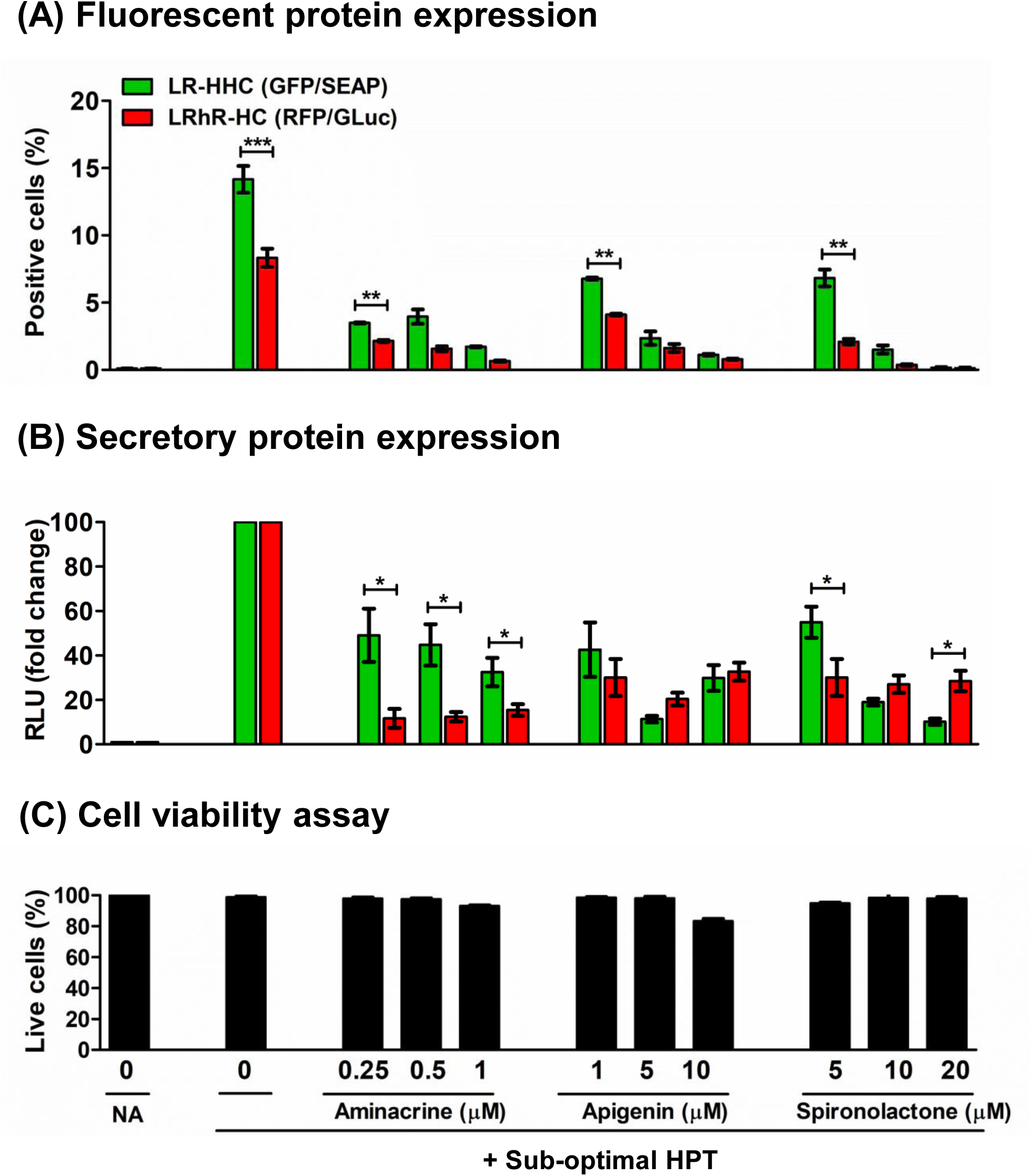
Latency-promoting agents can inhibit reporter gene expression from both LTRs. HIV-1 reporter gene expression from the LTRs was quantified following the treatment of the cells with varying concentrations of three known inhibitors of viral transcription. Note that the cells were also exposed to a sub-optimal HPT cocktail (HMBA 1.25 mM, PMA 1.25 ng/ml, and TNF-α 2.5 ng/ml) for 24 hrs. The expression of (**A**) fluorescent proteins and (**B**) secretory enzymes was analyzed. (**C**) Cell viability was monitored in all conditions. Data plotted in triplicate, mean ± SD, statistical test – Student’s t-test, *p < 0.05, **p < 0.01, ***p < 0.001.

### The dual-reporter cells offer a robust means for an HTS assay

An HTS assay must demonstrate a robust dynamic range and signal variability to distinguish between positive and cytotoxic agents. In an HTS assay, the Z’-score is a widely used and dimensionless statistical parameter (typically ranging from 0 to 1) that quantifies the separation and reproducibility between positive and negative controls (46). The Z′-score is the gold standard for evaluating assay quality before large-scale screening. To this end, we estimated the Z’-score values for both latency-reversal and latency-promoting assay formats of the dual reporter cells using the secretory proteins SEAP and GLuc.

The assays were performed in clear-bottom white 96-well plates. Each well contained 0.1 million cells, seeded in 100 µl of culture medium. The assay plates were used directly to measure GLuc and SEAP activities in the culture supernatant (as described in the materials and methods section). Notably, for the latency-reversal assay, the cells were pre-exposed to either a sub-optimal HPT activator or an optimal HPT activator cocktail (**Figure 5A**). In contrast, for the latency-promoting assay, the cells were treated with 10 µM Spironolactone in the presence of sub-optimal HPT activation (**Figure 5B**). The assays demonstrated Z’-scores in the ‘good to excellent’ range. For instance, the latency-reversal assay yielded Z’-scores of 0.7 and 0.72 for the SEAP and GLuc reporter enzymes, respectively, when the cells were treated with an optimal HPT cocktail of activators (**Figure 5A**). Similarly, the latency-promoting assay yielded Z’-scores of 0.5 and 0.76, respectively, for the two enzymes (**Figure 5B**). Thus, the dual-reporter cell line was found to be robust for screening LMAs.

**Figure 5.**
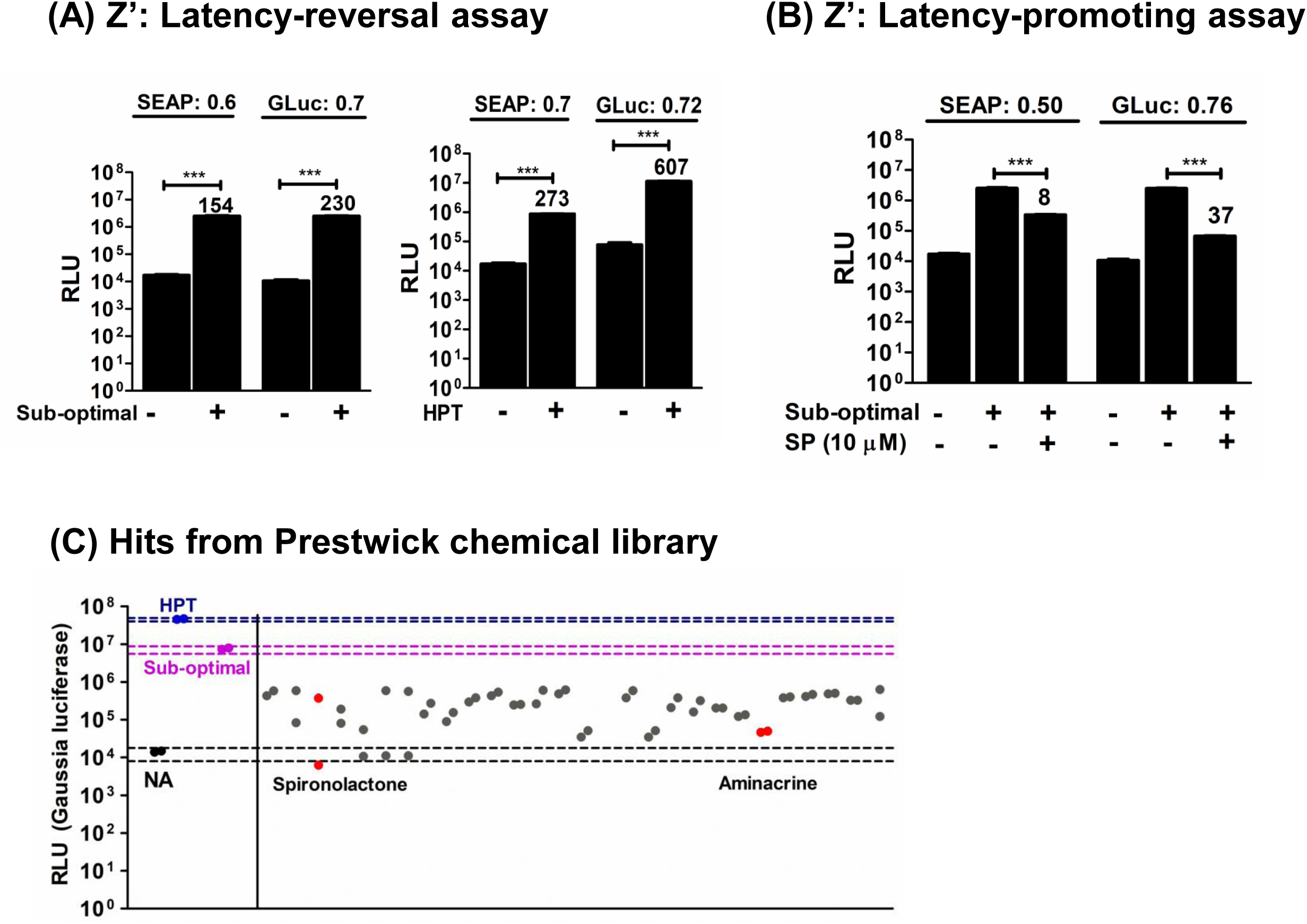
Dual-reporter Jurkat cell line is HTS compatible. Gaussia luciferase and secretory alkaline phosphatase encoded by the tough-to-activate LRhR-HC-LTR and the simple-to-activate LR-HHC-LTR, respectively, were used as the readout of the assay. Z’-score values for the assays are presented. Z-score values for (**A**) Latency-reversal assay at two different activation conditions, sub-optimal (sub-optimal, left) and HPT (right); and for (**B**) Latency-promoting assay was performed at a sub-optimal activation with 10 µM Spironolactone (SP). Fold change values are depicted. Data are plotted as n = 10, mean ± SD, statistics – Mann-Whitney U test, ***p < 0.001. (**C**) The Prestwick chemical library was screened to identify potential latency-promoting agents using the Gaussia luciferase assay. As shown, 27 hits are plotted as duplicates (grey circles) along with the assay controls: no-activation (black), positive-I (pink circles), and positive-II (blue circles); the dashed lines represent mean ± 3SD of the respective control.

### Screening of the Prestwick chemical library small molecules

Next, using the optimized HTS assay, we screened the PCL of 1,520 FDA-approved small molecules to identify potential LPAs. The assay format broadly followed the experimental procedure described above for determining Z’-scores. Each small molecule in the library was tested at 10 µM, in two wells, each on a separate plate, but at the same location. Necessary controls were included on all plates, including the optimal, sub-optimal HPT activator cocktail and the ‘no activation’ control. The levels of GLuc, encoded by the tough-to-activate LRhR-HC-LTR, were quantified in the medium at 24 hours.

Following testing the 1,520 SMs of the library in the assay, we identified approximately 200 potential hits that showed considerably lower GLuc activity than the positive controls. Considering the cytotoxicity, statistical significance, and discordant results between the duplicate wells, we narrowed the selected SM subset to 27 molecules (**Figure 5C)**. The subset of small molecules also contained Spironolactone and Aminacrine, previously identified to function as LPAs (43, 45), thus confirming the assay efficiency. Several other molecules identified here represent novel LPAs and warrant additional characterization using a range of HIV-1 infection models.

Further, we also screened a set of ten small molecules synthesised by Aurigene Oncology Ltd. to identify potential LMAs. We tested these molecules for their demonstrated inhibitory activity against cyclin-dependent kinases and/or as blockers of cell surface receptors regulating activation signals. Several host factors, including cellular kinases, potentiate Tat-mediated LTR transactivation. Preliminary screening was performed similarly to PCL screening, using GLuc as the primary readout and varying SM concentrations, in duplicate (**Figure 6A**). This screening identified four potential LPAs: CPD 1, 2, 4, and 8. These molecules demonstrated a dose-dependent inhibition of the LRhR-HC-LTR, as measured by GLuc activity (**Figure 6A**). While screening for LPAs, we also identified two potential LRAs. Since cells were also exposed to a sub-optimal activation, a compound exhibiting LRA activity would demonstrate increased GLuc expression compared to this activation control. This assay identified two such LRAs: CPD 6 and 9 (**Figure 6A**).

**Figure 6.**
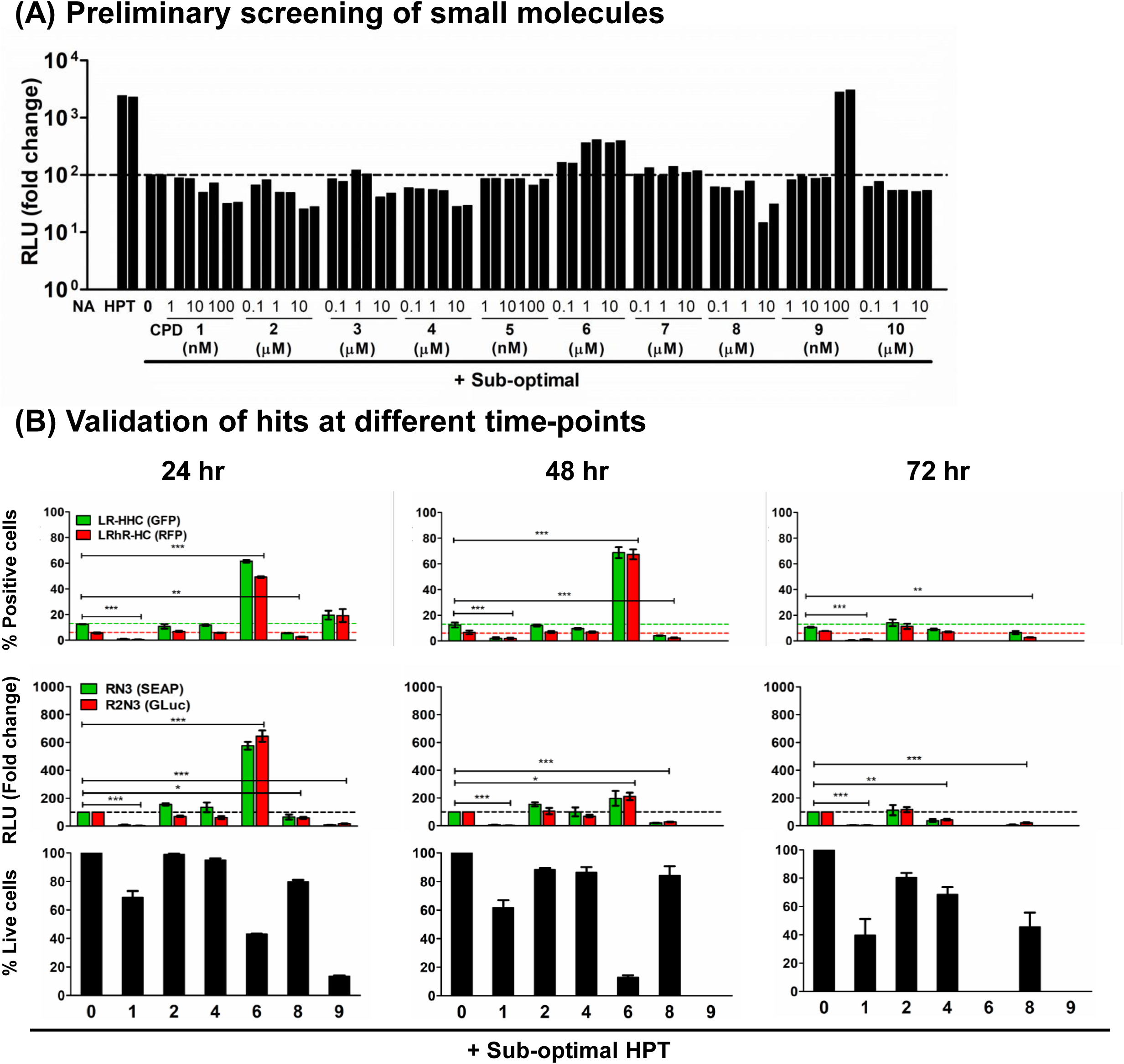
Screening of small molecules against HIV-1 gene expression. (**A**) A set of 10 small molecules were screened for latency modulation, each SM in duplicates. In the preliminary screening, six potential hits were identified - two as LRAs (CPD 6 and 9) and four as LPAs (CPD 1, 2, 4, and 8). In a modified method for LRA screening, molecules were incubated with a sub-optimal concentration of activators, and Gaussia luciferase values above sub-optimal activation control were considered as potential hits. (**B**) Hits (CPD 1, 2, 4, 6, 8, and 9) from preliminary screening were validated at three different time points. The expression of fluorescent (top panel) and secretory proteins (middle panel) were measured, along with cell cytotoxicity (bottom panel). This secondary screening validated the two most potent latency modulating agents, LPA: CPD 1 and LRA: CPD 6. Data plotted in triplicate, mean ± SD, comparison made against no activation control (0), statistical test – student’s t-test, *p < 0.05, **p < 0.01, ***p < 0.001.

The six hits from preliminary screening were validated at three different time points to assess their prolonged effect on viral transcription. These SMs were tested at their respective highest concentrations used in the initial analysis (**Figure 6A**), along with a sub-optimal concentration of cellular activators. The activities of both reporter proteins – secretory and fluorescent proteins were measured, along with cell cytotoxicity. The cell supernatant was collected every 24 hours, and the cells were fed fresh media containing sub-optimal activators and SMs. In the secondary assay, three SMs (CPD 1, 6, and 8) out of six showed consistent latency-modulating properties (**Figure 6B**).

The small molecule CPD 6 showed a potential latency-reversal activity, detectable only in the modified latency-reversal assay, in which cells are also exposed to sub-optimal activation conditions. However, beyond 24 hours, CPD 6 was highly toxic to the cells (**Figure 6B**, bottom panels). The secondary assay also validated two potential LPAs, CPD 1 and CPD 8. Both molecules showed reduced reporter activity at all three time-points, with CPD 1 being the most potent inhibitor. For instance, at 24 hours, CPD 1 reduced EGFP^+ve^ cells from 12.1 ± 0.5% to 1.5 ± 0.3%; and RFP^+ve^ cells from 5.1 ± 0.33% to 0.7 ± 0.1%. Likewise, CPD 8 reduced EGFP^+ve^ cells to 5.5 ± 0.56% and RFP^+ve^ cells to 2.5 ± 0.41% (**Figure 6B**, top panels). Comparable results were obtained for the secretory proteins (**Figure 6B**, middle panels). Compared to other compounds, CPD 1 and CPD 8 were less toxic beyond 24 hours of treatment. Furthermore, CPD 2 and CPD 4 exhibited no LMA activity, whereas CPD 9 was highly toxic to cells even at 24 hours. In summary, our analysis identified two potential LMAs, CPD 1 and CPD 6.

## III. Discussion

Combination antiretroviral therapy (cART) has substantially improved life expectancy in people living with HIV (PLWH). However, despite effective suppression of viral replication, ART does not fully eliminate low-level viral reactivation, often below detection thresholds. Persistent residual transcription contributes to chronic immune activation and associated comorbidities, including metabolic, renal, cardiovascular, and bone disorders etc (47). Because ART cannot eradicate latent reservoirs, lifelong treatment is required, which may promote the emergence of multidrug-resistant variants. These limitations highlight the need for a new class of antiviral agents, latency-modulating agents (LMAs), that directly target the HIV-1 MTRC rather than viral proteins.

The remarkable stability of HIV latent reservoirs remains a major barrier to eradication or functional cure (48, 49). The shock-and-kill strategy has shown limited success, likely because latency is maintained through multiple regulatory layers (50). Consequently, complete reactivation and clearance of all replication-competent proviruses may not be practically achievable using this approach.

In contrast, the block-and-lock strategy offers several conceptual and technical advantages. Unlike shock-and-kill, which requires global activation of latent reservoirs, block-and-lock aims only to suppress spontaneous viral transcription(51, 52). This approach is more compatible with small-molecule LPAs and may avoid inflammatory toxicities associated with reservoir activation. Notably, a small molecule, didehydro-cortistatin A (dCA), derived from a sea sponge and known to inhibit Tat-mediated transactivation, has been shown to suppress residual HIV reactivation (53). Unfortunately, dCA is a chemically complex molecule that is not amenable to chemical modification and large-scale synthesis, despite its proven specificity in suppressing HIV transcription. These limitations underscore the need to identify new LPAs, necessitating robust reporter-based HTS platforms.

Most conventional HTS assays use the canonical HIV-1 LTR, but intrinsic features make it poorly suited for screening transcription-specific inhibitors (54). Tat-independent transcription is heavily dependent on NF-κB signaling (31, 55), and both Tat-dependent and -independent transcription rely entirely on host machinery, leading to poor viral specificity (32, 56). Consequently, many hits target general host transcription pathways rather than the viral promoter. For example, JTK-101, an SMI, potently suppresses both TNF-α-induced and Tat-mediated LTR transactivation at nanomolar concentrations by inhibiting the cellular P-TEFb complex (57). Similarly, Ro 5-3335, another anti-Tat SMI, strongly inhibits Tat-dependent transactivation and is thought to exert its anti-HIV activity by suppressing a host factor required for Tat function (58). Furthermore, the canonical LTR also produces unusually high transcriptional noise compared with cellular promoters (34, 35), a feature thought to aid transitions between active and latent states (36), but one that elevates background and increases false positives in HTS. Thus, an LTR that is relatively more resistant to the cellular transcriptional noise is expected to be more suited for developing an HTS assay for HIV.

An effective LPA should selectively suppress viral transcription, exert durable antiviral effects, and function alongside ART. These criteria guided the development of the HTS assay described here. HIV-1C LTR variants with duplicated transcription factor binding sites (TFBS), particularly those with an additional RBEIII motif and no corresponding NF-κB duplication, exhibit elevated activation thresholds (23). The LRhR-HC-LTR used in this study shows strong resistance to basal transcriptional noise,(25) making it particularly well-suited for HTS while minimizing false-positive hits.

Compounds identified through Prestwick Chemical Library screening suggest the involvement of host factors distinct from the canonical transcription machinery in regulating HIV transcription. For example, spironolactone suppresses Tat-dependent transcription by promoting degradation of XPB, a TFIIH component, without broadly destabilizing general transcription factors (45). Aminacrine inhibits viral transcription indirectly by inducing p21WAF1, which sequesters cyclin T1/CDK9 away from the HIV LTR (59). Additional hits from our screen may represent novel LMAs, although further validation is required to establish specificity and mechanism.

The latency-resistant phenotype of the LRhR-HC-LTR can be explained by its TFBS composition, which governs transcriptional output and ON/OFF switching. HIV-1 LTRs recruit activating and repressing host complexes, including NF-κB, NFAT, AP-1, RBF-2, and Sp1, whose effects depend on cellular activation state(60–62). While NF-κB sites promote transcription, the upstream RBEIII motif imposes repression under resting conditions (66, 67). Our previous findings show that adding an extra RBEIII motif without co-duplicating NF-κB sites (e.g., LRhR-HC-LTR), and even the possible absence of one functional NF-κB motif, designated as ‘h’ here, markedly suppresses transcription across multiple viral systems and increases resistance to latency reversal (24, 25). Patient-derived CD4^⁺^ T cell reactivation assays further confirm the biological relevance of this suppressive phenotype.

The dual-reporter cell line described here offers several advantages over existing models (68–70). It enables parallel readouts from a highly stringent LTR (LRhR-HC) and a canonical LTR (LR-HHC), balancing specificity with screening flexibility. The system supports both fluorescent and secreted enzyme reporters, allowing non-destructive, multiplexed quantification. Depending on activation conditions, the same platform can screen for LPAs or LRAs. High Z′-scores demonstrate excellent assay robustness, sensitivity, and reproducibility. Importantly, the system isolates the viral MTRC by including only the LTR and Tat, minimizing confounding effects from other viral elements, unlike a few previously reported screening systems (63–70). Finally, this is the first reporter cell line derived from an HIV-1C backbone. Despite this, it responds comparably to standard LRAs and LPAs used in HIV-1B systems, suggesting broad subtype applicability.

In summary, we developed a dual-reporter HIV-1C–based cell line that combines high assay stringency, resistance to transcriptional noise, and strong reproducibility, providing a versatile HTS platform for identifying both latency-promoting and latency-reversing agents.

## IV. Materials and methods

### Construction of viral vectors

The LRhR-HC-LTR-GLuc-IRES-Tat-mScarlet vector was generated from a previously constructed LRhR-HC-LTR-d2EGFP-IRES-Tat-mScarlet backbone (unpublished). Gaussia luciferase (GLuc) fused to the FKBP12 degradation domain was generated using an overlap-PCR strategy. The GLuc fragment was amplified from a previously reported Tat- and Rev-dependent construct, pCL-dmScarlet-GLuc-HHC (25), using primers N4328 and N4329 (See Table 1). The FKBP fragment was amplified from a previously described pcLdGITRD vector (80) using primers N4248 and N4249. The final GLuc-FKBP amplicon was produced by overlap-PCR using the two amplified fragments and cloned into the LRhR-HC-LTR-d2EGFP-IRES-Tat-mScarlet vector by replacing d2EGFP using BamHI (#R0136S, New England Biolabs) and EcoRI (#R0101S, New England Biolabs) restriction sites.

**Table 1.**
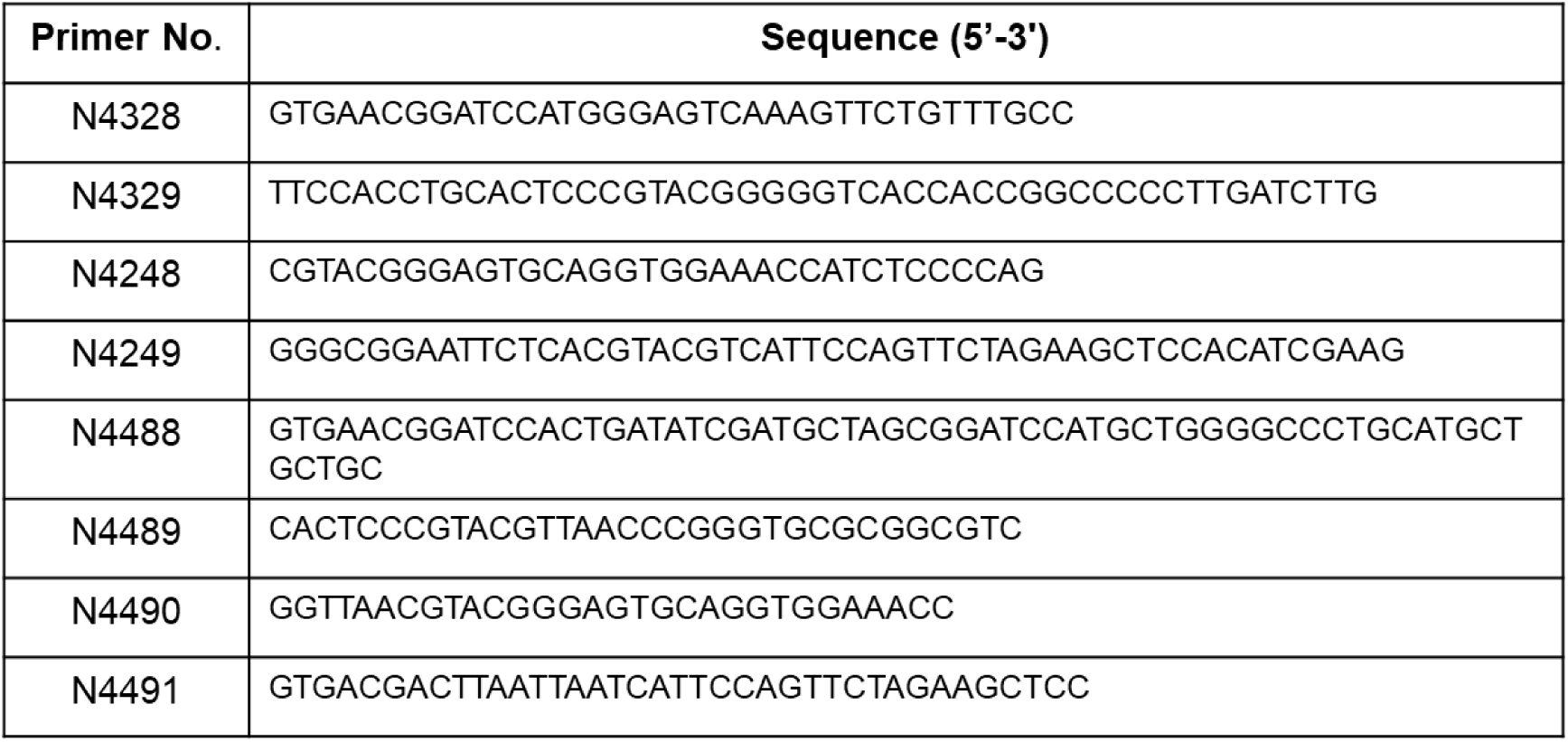
Primer Sequences.

The LR-HHC-LTR-SEAP-IRES-EGFP vector was constructed from a previously reported LR-HHC-LTR-d2mScarlet-IRES-EGFP vector (25). SEAP was amplified from pLTRD24-SEAP-IRES-EGFP (#ARP-11788, NIH Reagents Program) using primers N4488 and N4489. The FKBP fragment was amplified from the pcLdGITRD vector (80) using primers N4490 and N4491. The SEAP-FKBP fusion fragment was generated by overlap-PCR and cloned into the pCL-d2mScarlet-IRES-EGFP vector (25) by replacing d2mScarlet using EcoRV (#R0101S, New England Biolabs) and PacI (#R0547S, New England Biolabs) restriction sites. Primer sequences are listed in **Table 1**.

### Cell culture

HEK293T cells (#CRL-3216, ATCC) were maintained in DMEM (AL007-500ML, VASA Scientific, India) supplemented with 10% fetal bovine serum (FBS; #S-FBS-US-025, Life Technologies), and GPS – 2 mM glutamine (#G8540, Sigma-Aldrich), 100 units/l penicillin G (#P3032, Sigma-Aldrich), and 100 μg/ml streptomycin (#S9137, Sigma-Aldrich). Jurkat E6-1 cells (#ARP-177, NIH Reagents Program) were cultured in RPMI 1640 medium (#R4130, Sigma-Aldrich) supplemented with 10% FBS and GPS. All cells were maintained at 37°C in a humidified incubator with 5% CO₂.

### Preparation of viral stocks

Viral stocks were generated in HEK293T cells by co-transfecting individual viral vectors with third-generation lentiviral packaging plasmids using the calcium phosphate transfection method (81). HEK293T cells were cultured in DMEM supplemented with 10% FBS and GPS. For each transfection in a 6-well plate at 40–50% confluency, a plasmid cocktail consisting of 2 µg viral vector, 1 µg psPAX2 (#11348, NIH AIDS Reagent Program), 0.65 µg pHEF-VSVG (#4693, NIH AIDS Reagent Program), and 0.35 µg pCMV-Rev (#1443, NIH AIDS Reagent Program) was used. Six hours post-transfection, the medium was replaced with fresh DMEM containing 10% FBS and GPS. Forty-eight hours later, viral supernatants were harvested, clarified by centrifugation at 1,500 rpm for 5 minutes, aliquoted, and stored at −80°C until use.

### Generation of a dual-reporter cell line

The dual-reporter cell line was generated through a two-step infection and selection process. Initially, Jurkat cells were infected with a viral strain harboring the LRhR-HC-LTR at approximately 10% infectivity. Infected cells were activated using an HPT cocktail consisting of 5 mM HMBA (Sigma #224235), 5 ng/ml PMA (Sigma #P1585), and 10 ng/ml TNF-α (Miltenyi Biotech #130-094-019), followed by single-cell sorting into 96-well plates (Biolite™, #130188). Each well contained 100 µl of fresh RPMI medium (Invitrogen #3821000) supplemented with 10% FBS and 100 µl of conditioned medium from Jurkat cells, along with GPS components. Clonal populations were screened for GLuc and RFP expression under activating conditions (data not shown). A single clone exhibiting the highest reporter activity was selected and subsequently co-infected with the second viral vector, LR-HHC, using the same experimental strategy. Following HPT activation, cells expressing both EGFP and RFP were single-cell sorted into 96-well plates and expanded. Four clonal sub-populations were selected for further characterization.

### Characterization of a dual-reporter cell line

Four clonal cell lines (clones 9, 14, 19, and 39) were characterized by activating 0.2 million cells in 200 µl RPMI supplemented with 10% FBS and GPS for 24 hours using different combinations of cellular activators. Cells were treated with either sub-optimal HPT conditions (0.1 or 1 mM HMBA, 1.5 ng/ml PMA, and 2.5 ng/ml TNF-α) or optimal HPT conditions (5 mM HMBA, 5 ng/ml PMA, and 10 ng/ml TNF-α). Twenty-four hours post-activation, reporter enzyme activity was quantified from harvested cells and spent culture supernatants. For flow cytometric analysis, cells were stained with a live/dead fixable far-red dead cell stain (Thermo Scientific #L34973) diluted 1:10,000 in PBS. Cells were washed once with PBS, pelleted at 800 g for 5 minutes at room temperature, and then resuspended in 100 µl of diluted stain. The cells were incubated at 4°C for 30 minutes. After two additional PBS washes, the cells were resuspended in 200 µL PBS and analysed on a BD Aria III flow cytometer by acquiring 10,000 events per sample. Data analysis was performed using FCS Express 6. To measure GLuc and SEAP activity in the culture supernatant, assay plates were placed directly on a multimode plate reader (Thermo Scientific Varioskan^TM^ LUX, #3020-973). Commercial kits were used to quantify reporter enzyme activities: GLuc using the Pierce Gaussia Luciferase Glow Assay Kit (Thermo Scientific, #16161) and SEAP using the Secreted Alkaline Phosphatase Reporter Gene Assay Kit (Cayman, #600260), both according to the manufacturer’s protocols.

### Latency modulating agents

Five standard LRAs (Ingenol-3-angelate, JQ1, Prostratin, Romidepsin, and SAHA) and three LPAs (Aminacrine, Apigenin, and Spironolactone) were evaluated. A total of 0.2 million cells were treated with varying concentrations of LPAs in 200 µl RPMI supplemented with 10% FBS and GPS in a 48-well plate for 24 hours. For testing the LRAs, cells were incubated with varying concentrations of LRAs, along with a ‘No activation’ as a negative control, containing only 0.1% DMSO. For testing the LPAs, cells were incubated with varying concentrations of the modulators in the presence of a sub-optimal HPT activator cocktail (1.25 mM HMBA, 1.25 ng/mL PMA, and 2.5 ng/mL TNF-α) for 24 hours. Following the treatment, the levels of the reporter enzymes in the spent medium samples were quantified as described above. Cells were stained with a live/dead stain and analyzed by flow cytometry as mentioned above. To measure GLuc and SEAP activity in the culture supernatant, assay plates were placed directly on a multimode plate reader (Thermo Scientific Varioskan^TM^ LUX, #3020-973). Commercial kits were used to quantify reporter enzyme activities: GLuc using the Pierce Gaussia Luciferase FLASH Assay Kit (Thermo Scientific, #16158) and SEAP using the Secreted Alkaline Phosphatase Reporter Gene Assay Kit (Cayman, #600260), both according to the manufacturer’s protocols.

### Determination of the Z’-score

The assay was configured in clear-bottom white 96-well plates (Thermo Scientific #165306), with each well containing 0.1 million reporter cells in 100 µl RPMI supplemented with 10% FBS. Peripheral wells were excluded from analysis and filled with PBS to minimize evaporation. To calculate the Z’-score for the latency-reversal assay, cells were activated for 24 hours using either optimal (5 mM HMBA, 5 ng/ml PMA, 10 ng/ml TNF-α) or sub-optimal (1.25 mM HMBA, 1.25 ng/ml PMA, 2.5 ng/ml TNF-α) HPT conditions. GLuc and SEAP activities were quantified directly from culture supernatants using a multimode plate reader (Thermo Scientific Varioskan™ LUX, #3020-973). GLuc activity was measured using the Pierce™ Gaussia Luciferase FLASH Assay Kit (#16158, Thermo Scientific), and SEAP activity was quantified using the Secreted Alkaline Phosphatase Reporter Gene Assay Kit (#600260, Cayman), following the manufacturers’ protocols. For latency-promoting assays, cells were treated with 10 µM Spironolactone in the presence of sub-optimal HPT activation. Sub-optimal activation alone served as the positive control, and no activation served as the negative control.

### Screening of a small molecule library

Small molecules from the Prestwick Chemical Library (PCL) (ver.19 ref.PCL-10-100-P96, Prestwick Chemical, France) were diluted in DMSO to a working concentration of 100 µM and dispensed into white 96-well plates (Thermo Scientific #136101). Plates were sealed and stored at −80°C until use. Ten additional compounds from Aurigene Limited (CPD 1 to 10) were dissolved in DMSO at 10 mM, and 2 mM working stocks were prepared in RPMI containing 20% DMSO and stored at −20°C.

Primary screening of the PCL was conducted in clear-bottom white 96-well plates using 0.1 million cells per well in 100 µl RPMI supplemented with 10% FBS. GLuc activity was measured as described above for Z’-score determination. Peripheral wells were excluded from analysis. Each compound was tested at 10 µM in duplicate wells across separate plates, with wells positioned identically on each plate. Each plate included no-activation, sub-optimal-activation, and optimal-activation controls in duplicate. For latency-promoting assays, compounds were tested at 10 µM in the presence of sub-optimal HPT activation.

Compounds from Aurigene Oncology Ltd. underwent a sequential screening strategy. Primary LSA screening was performed similarly to the PCL screening but tested at three different concentrations. Validation involved measuring all four reporter outputs: GLuc, SEAP, EGFP, and mScarlet. These assays were conducted in 48-well plates containing 0.2 million cells per well in 200 µl RPMI supplemented with 10% FBS. After 24 hours, culture supernatants were collected for GLuc and SEAP measurements, and cells were stained with a live/dead dye and analyzed by flow cytometry to quantify fluorescent reporter expression and cell viability.

### Statistical analysis

All data plotting and statistical analyses were performed using GraphPad Prism software (version 5.02). Luciferase data acquisition was carried out using SkanIt RE software (version 7.02). The specific statistical tests used for each experiment, along with the corresponding p-values, are indicated in the relevant figure legends. Flow cytometry data were analyzed using FCS Express 6 software (De Novo Software, Los Angeles, CA).

## Author contributions

C.S. performed research, analyzed data, and wrote the paper. A.R.C., S.G., and Y.G. performed research and analyzed data. A.P. provided inputs in statistical analysis and manuscript editing. P.D., S.T., M.R., and S.S. designed small molecules CPD 1 to 10. R.M. provided the Prestwick Chemical Library, contributed to the study design, and manuscript editing. U.R. designed the research and wrote the paper.

## Acknowledgment

We thank Dr. Prasanta K Dash, Department of Pharmacology and Experimental Neuroscience, University of Nebraska Medical Centre, Omaha, Nebraska, USA, for providing inputs in manuscript editing.

